## Supplementary Figures and Tables for "Neuronal NPR-15 modulates molecular and behavioral immune responses via the amphid sensory neuron-intestinal axis in *C. elegans*": Otarigho et al_Supplementary Figures.docx

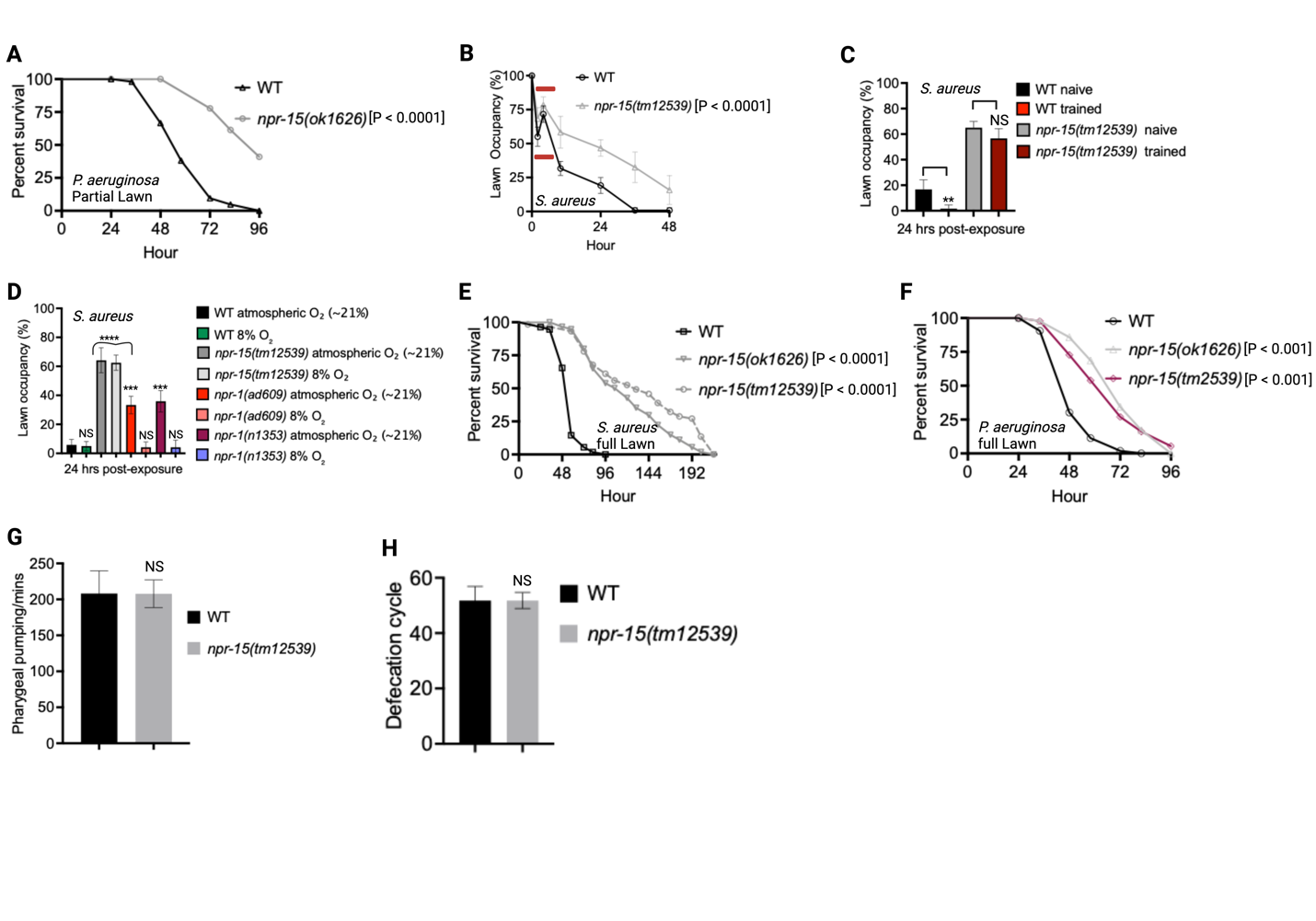

**Figure S1. NPR-15 loss-of-function exhibited pathogen resistance independent of brood size and oxygen-independent avoidance behavior. Related to Figure 1.**

A. Wild type (WT) and *npr-15(ok1626)* animals were exposed to *P. aeruginosa* partial lawn and scored for survival.
B. Lawn occupancy of WT and *npr-15(tm12539)* animals exposed to a partial lawn of *S. aureus*. *** p < 0.001. The marked region shows re-occupancy.
C. Lawn occupancy of naïve and trained WT and *npr-15(tm12539)* animals exposed to a partial lawn of *S. aureus*. ** p< 0.001 **** p < 0.0001.
D. Lawn occupancy of WT, *npr-15(tm12539), npr-1(ad609)*, and *npr-1(n1353)* animals exposed to a partial lawn of *S. aureus* at 24 hours in 8% and atmospheric oxygen (~21%). Bars represent means while error bars indicate SD; ****p < 0.0001, *** p < 0.001, and NS = Not Significant.
E. WT, *npr-15(tm12539),* and *npr-15(ok1626)* animals were exposed to *S. aureus* full lawn and scored for survival.
F. WT, *npr-15(tm12539),* and *npr-15(ok1626)* animals were exposed to *P. aeruginosa* full lawn and scored for survival.
G. The pharyngeal pumping rate of WT and *npr-15(tm12539)* animals. Bars represent means while error bars indicate SD; P =NS.
H. Defecation cycle of WT and *npr-15(tm12539)* animals*­­*­. Bars represent means while error bars indicate SD; P =NS.

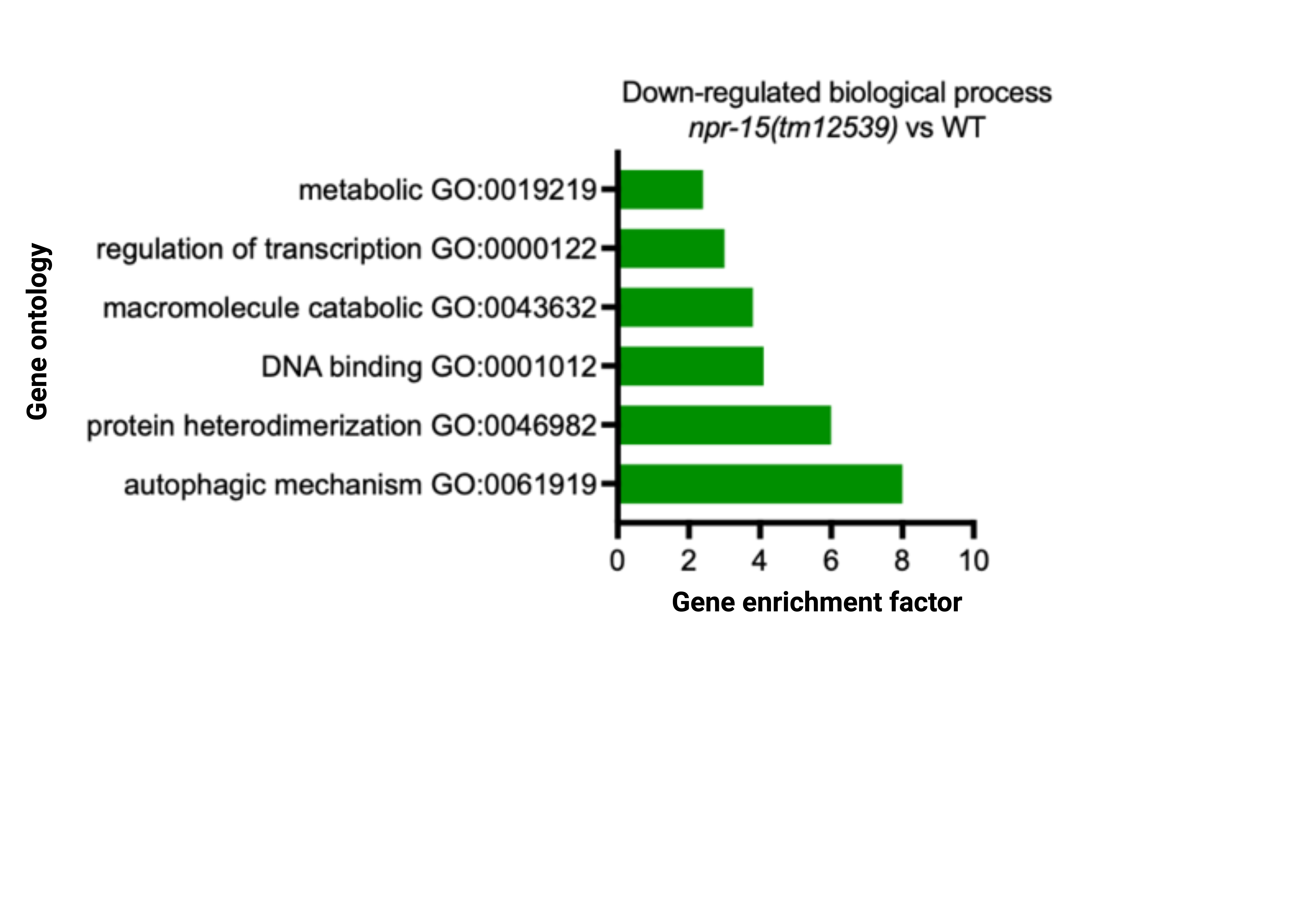

**Figure S2. Downregulated biological process and upregulated immune pathways/gene number in *npr-15(tm12539)* animals. Related to Figure 2.** Gene ontology analysis of downregulated genes in *npr-15(tm12539)* vs. WT animals. The result was filtered based on significantly enriched terms, with a Q value < 0.1.

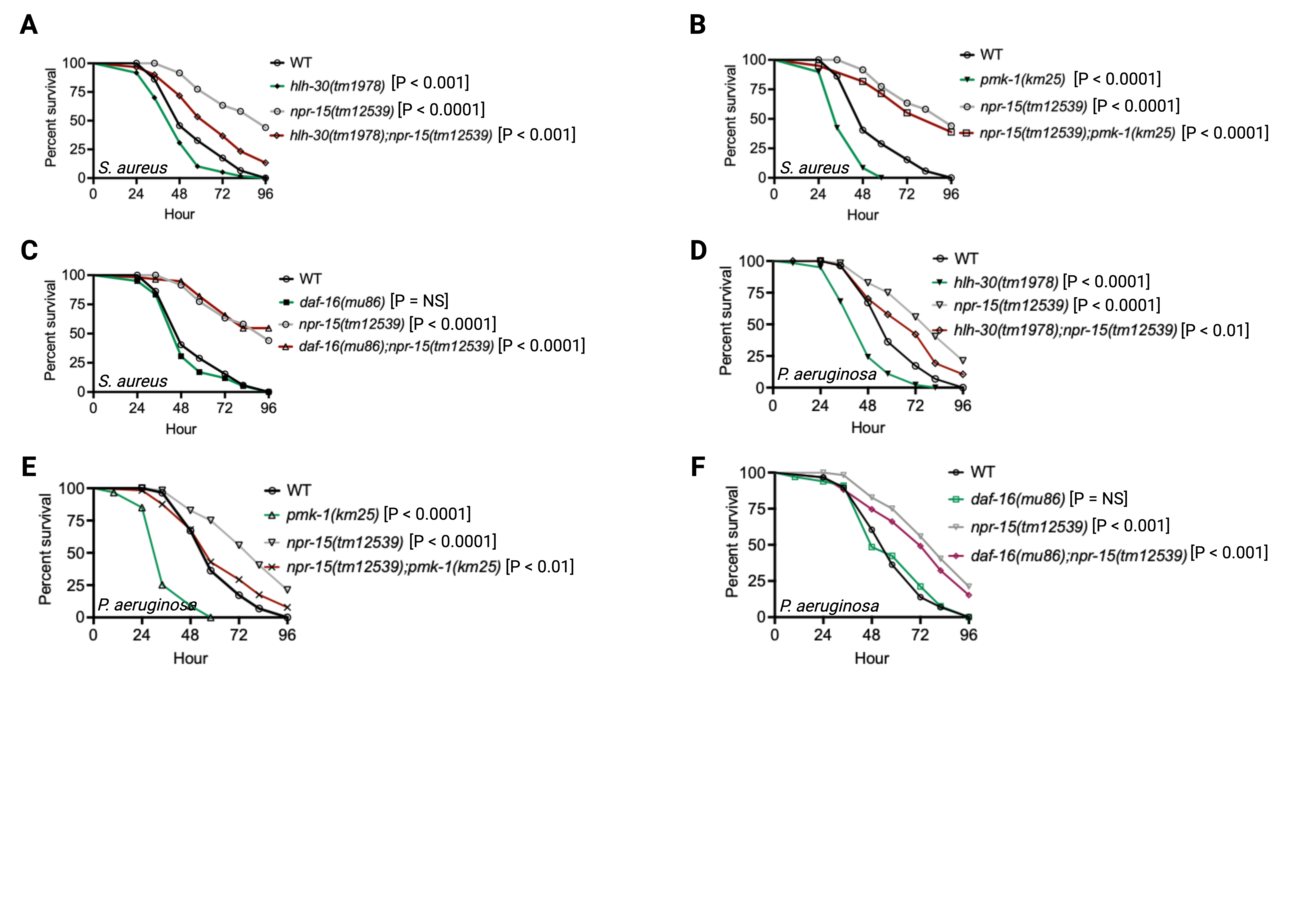

**Figure S3. NPR-15 loss-of-function enhances immunity via ELT-2 and HLH-30 when exposed to *S. aureus*. Related to Figure 3.**

A. WT, *hlh-30(tm1978),* *npr-15(tm12539),* and *hlh-30(tm1978);npr-15(tm12539)* animals were exposed to *S. aureus* full lawn and scored for survival.
B. WT, *pmk-1(km25),* *npr-15(tm12539)*, *npr-15(tm12539);pmk-1(km25)* animals were exposed to *S. aureus* full lawn and scored for survival.
C. WT, *daf-16(mu86)*, *npr-15(tm12539),* and *daf-16(mu86)*;*npr-15(tm12539)* animals were exposed to *S. aureus* full lawn and scored for survival.
D. WT, *hlh-30(tm1978),* *npr-15(tm12539),* and *hlh-30(tm1978);npr-15(tm12539)* animals were exposed to *P. aeruginosa* full lawn and scored for survival.
E. WT, *pmk-1(km25),* *npr-15(tm12539)*, *npr-15(tm12539);pmk-1(km25)* animals were exposed to *P. aeruginosa* full lawn and scored for survival.
F. WT, *daf-16(mu86)*, *npr-15(tm12539),* and *daf-16(mu86)*;*npr-15(tm12539)* animals were exposed to *P. aeruginosa* full lawn and scored for survival.

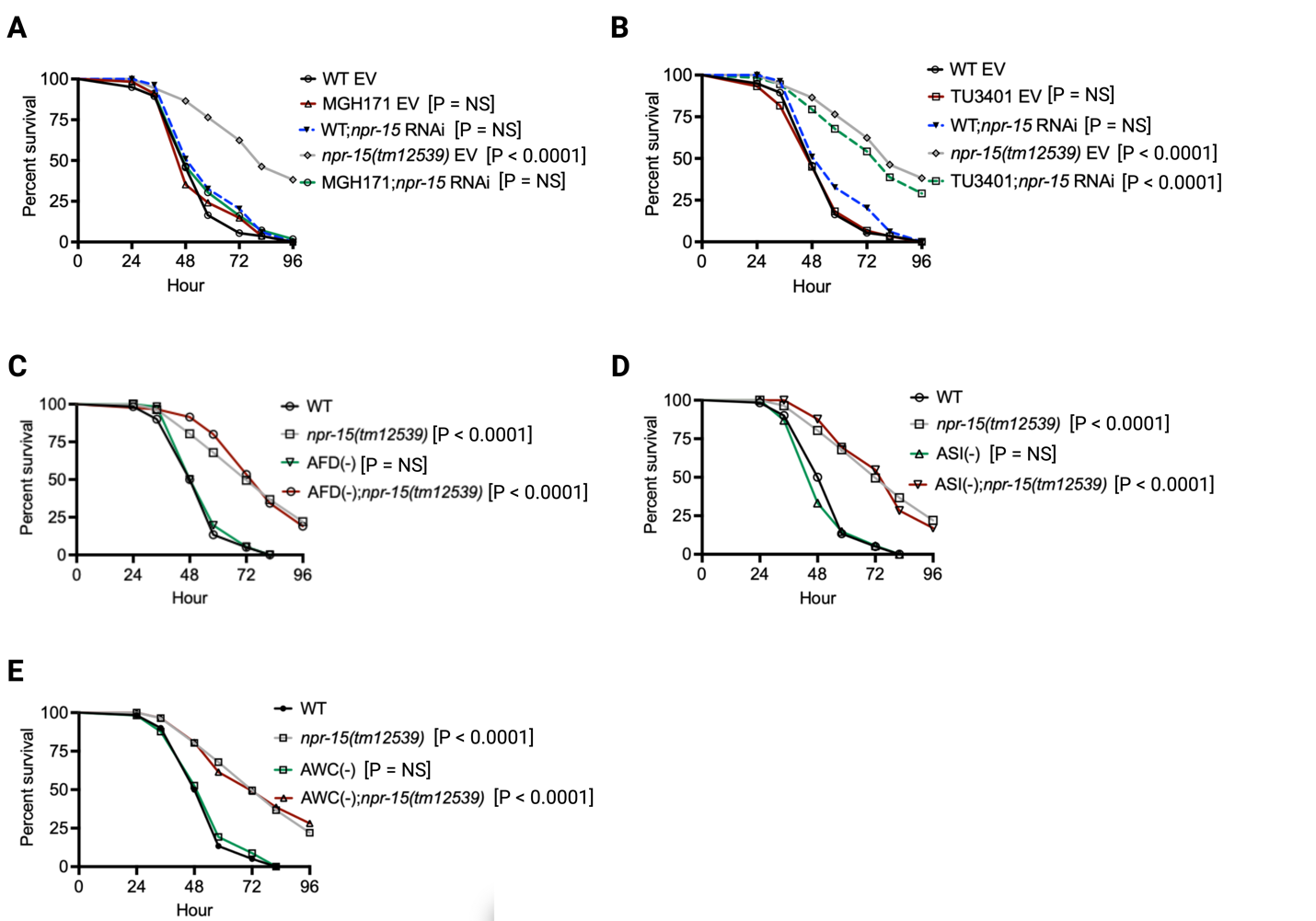

**Figure S4: Survival of other NPR-15-expressing neuron-ablated strains in *S. aureus* infection. Related to Figure 4.**

A. WT and MGH171 animals treated with *npr-15* RNAi, exposed to *S. aureus* full lawn alongside *npr-15(tm12539)* animals, and scored for survival. EV, empty vector RNAi control.
B. WT and TU3401 animals treated with *npr-15* RNAi, exposed to *S. aureus* full lawn alongside *npr-15(tm12539)* animals, and scored for survival. EV, empty vector RNAi control.
C. WT, *npr-15(tm12539),* AFD(-), and AFD(-);*npr-15(tm12539)* animals were exposed to *S. aureus* and scored for survival.
D. WT, *npr-15(tm12539),* ASI(-), and ASI(-);*npr-15(tm12539)* animals were exposed to *S. aureus* and scored for survival.
E. WT, *npr-15(tm12539),* AWC(-), and AWC(-);*npr-15(tm12539)* animals were exposed to *S. aureus* and scored for survival.

**Figure S5. NPR-15 controls avoidance behavior independent of immunity, neuropeptide pathways, and oxygen. Related to Figures 3 and 5.**

A. Lawn occupancy of WT and *npr-15(tm12539)* animals treated with *elt-2* RNAi before exposure to a partial lawn of *S. aureus* at 24 hours. Bars represent means while error bars indicate SD; ***p < 0.0001, and NS = Not Significant.
B. Lawn occupancy of WT and *npr-15(tm12539)* animals treated with *pmk-1* RNAi before exposure to a partial lawn of *S. aureus* at 24 hours. Bars represent means while error bars indicate SD; ***p < 0.0001, and NS = Not Significant.
C. Lawn occupancy of WT and *npr-15(tm12539)* animals treated with *daf-16* RNAi before exposure to a partial lawn of *S. aureus* at 24 hours. Bars represent means while error bars indicate SD; ***p < 0.001, and NS = Not Significant.
D. Lawn occupancy of WT and *npr-15(tm12539)* animals treated with *hlh-30* RNAi before exposure to a partial lawn of *S. aureus* at 24 hours. Bars represent means while error bars indicate SD; ***p < 0.001, and NS = Not Significant.
E. Lawn occupancy of WT and *npr-15(tm12539)* animals treated with different neuropeptide gene RNAi before exposure to a partial lawn of *S. aureus* at 24 hours. Bars represent means while error bars indicate SD; ****p < 0.0001, and NS = Not Significant.

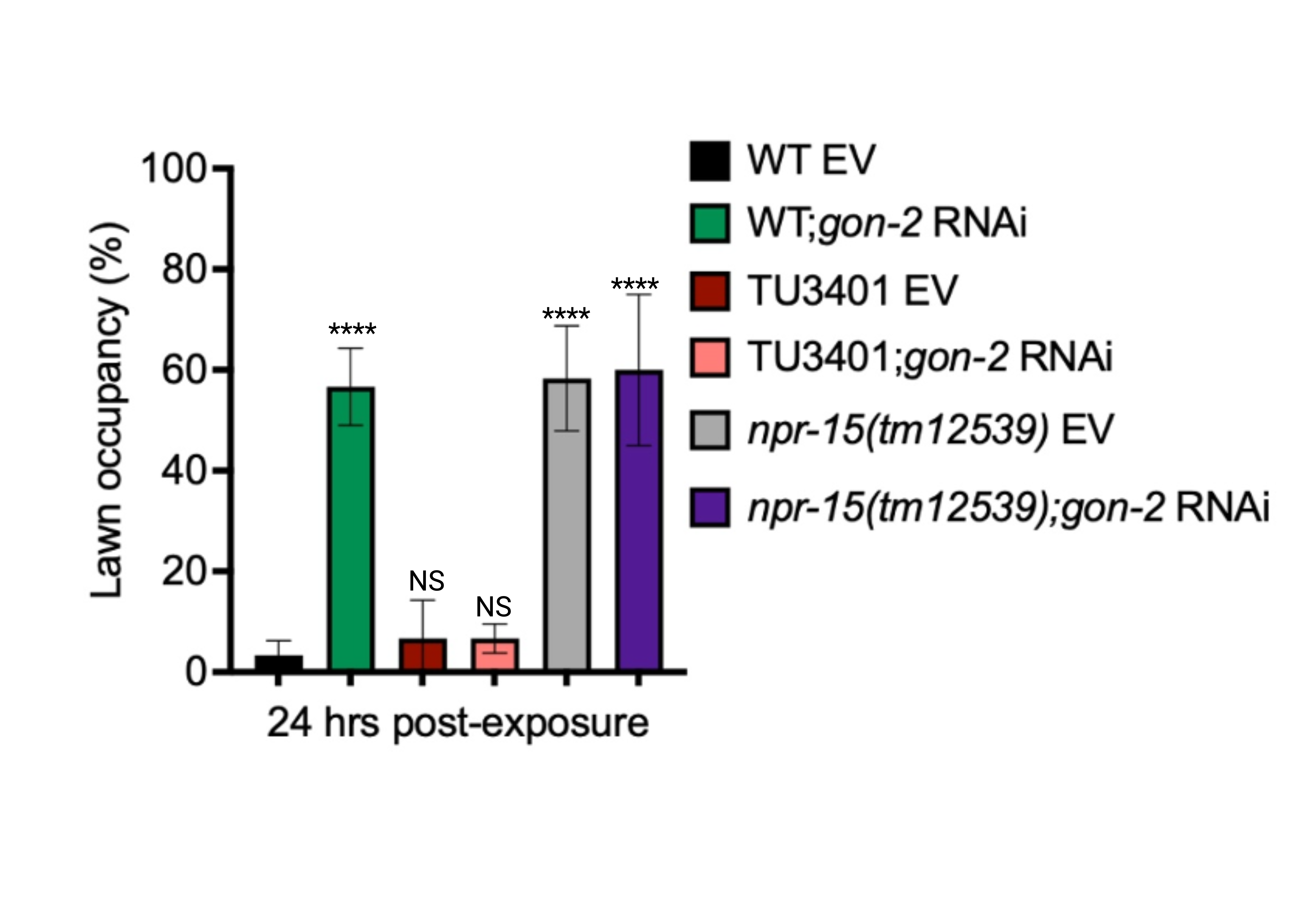

**Figure S6. The TRPM channel GON-2 control avoidance is independent of the nervous system. Related to Figure 5.** WT, TU3401, and *npr-15(tm12539)* fed with *gon-2* RNAi and were exposed to the partial lawn of *S. aureus* and scored for lawn occupancy. EV, empty vector RNAi control. Bars represent means while error bars indicate SD; ****p < 0.0001, and NS = Not Significant.
